# Unifying comprehensive genomics and transcriptomics in individual cells to illuminate oncogenic and drug resistance mechanisms

**DOI:** 10.1101/2022.04.29.489440

**Authors:** Jeffrey R. Marks, Jon S. Zawistowski, Isai Salas-González, Tia A. Tate, Tatiana V. Morozova, Jeff G. Blackinton, Durga M. Arvapalli, Swetha D. Velivela, Gary L. Harton, Charles Gawad, Victor J. Weigman, E. Shelley Hwang, Jay A.A. West

## Abstract

Discovering transcriptional variation in the absence of underlying genomic contributions hinders understanding of molecular mechanisms of disease. To assess this coordination in individual cells, we leveraged a new workflow, ResolveOME, exploiting the attributes of primary template-directed amplification (PTA) to enable accurate, complete-genome assessment of single-nucleotide variation in conjunction with full-transcript RNA-seq.

In cultured AML cells resistant to the FLT3 inhibitor quizartinib, we uncovered a *FLT3* missense mutation and matched transcript upregulation of AXL signal transduction and enhancer factor modulation driving resistance. In primary breast cancer cells, we detected oncogenic *PIK3CA* N345K mutations and heterogeneous classes of chromosomal loss and were empowered to interpret these genotypes with the crucial knowledge of cell identity and state derived from the transcriptome. The study reinforces the plasticity of the genome in conjunction with expected transcriptional modulation, leading to combinatorial alterations that affect cellular evolution that can be identified through application of this workflow to individual cells.

## INTRODUCTION

Remarkable variation between individual cells defines cell neighborhoods. A multitude of studies have described a myriad of tumorigenic mechanisms. However, cancer etiology remains driven by speculation, due to poor resolution of analyses that utilize population averages of cells to define heterogeneity. Borne out in the molecular complexity of single cells, the underlying resiliency of drug resistant cancer cells is driven by the convergence of single nucleotide variation (SNV), copy number variation (CNV) and transcriptional adaptation^1^. While one of these modes can be a dominant driver, they are not likely mutually exclusive, but rather, work synergistically to change cell state leading to progression or resistance^2^. To improve our understanding of heterogeneity, substantial improvements in assay resolution are required. Recent methods have pioneered simultaneously monitoring both RNA and DNA in single cells, however these produce uneven genome coverage and low allelic balance, limiting the ability to assess single nucleotide variation genome-wide with accuracy^3–5^.

To overcome this challenge, we enhanced the previously-characterized PTA^6^ workflow by extending it to a second modality of transcriptome enrichment. The PTA method is differentiated through superior genome coverage and uniformity that results in the highest reported precision and sensitivity at single-cell resolution^6–8^. The unified method layers full-transcript RNA quantification, delineating expression and isoform structure, onto the DNA analysis exposing CNV and SNV, allowing for the linkage of the genetic alterations and transcriptional programs at single cell resolution.

Prior multi-omic efforts that paired genomic and transcriptomic information from the same single cell have the primary shortcoming of incomplete genome coverage and associated non-uniformity of coverage, leaving uncovered genomic valleys that may harbor deleterious single nucleotide variants that would remain undetected. Indeed, prior whole genome amplification (WGA) methods^3–5^ are outperformed by PTA^6^ in terms of genomic coverage, allelic balance and SNV calling metrics. While clonal evolution at the SNV/CNV level in a primary patient sample has been accomplished by G&T-seq, the study was limited to a candidate gene survey defined by 59 oncogenes^9^. The workflow reported here builds upon the improved genome coverage and enhanced synced transcriptome performance, enabling novel linkage from the genome to the transcriptome which in several cases improved our understanding of the events that may impact patient outcome. We demonstrate the significance of these measurements, whereby single nucleotide variation fundamentally affects cell state^10^ and tumor progression^11,12^. The utility of these unified “omic” layers are highlighted with heterogenous genomic variation and consequential phenotypic alterations in single cells that both are correlated with the development of resistance to a targeted therapeutic in acute myeloid leukemia cells and in oncogenic mechanisms in primary breast cancer cells that could not be inferred by genomic or transcriptomic analyses in isolation.

## RESULTS

### Genomic and transcriptomic performance metrics

To benchmark this workflow (**Figure 1a**), the NIST Genome in a Bottle (GIAB) cell line, NA12878^13^ was used to define the technical performance by comparing the RNA or DNA chemistry in isolation to the combined workflow. Efficiency of the yield of the unified DNA and RNA protocol, along with other QC metrics are shown in **Supplementary Figure 1a**.

**Fig. 1.**
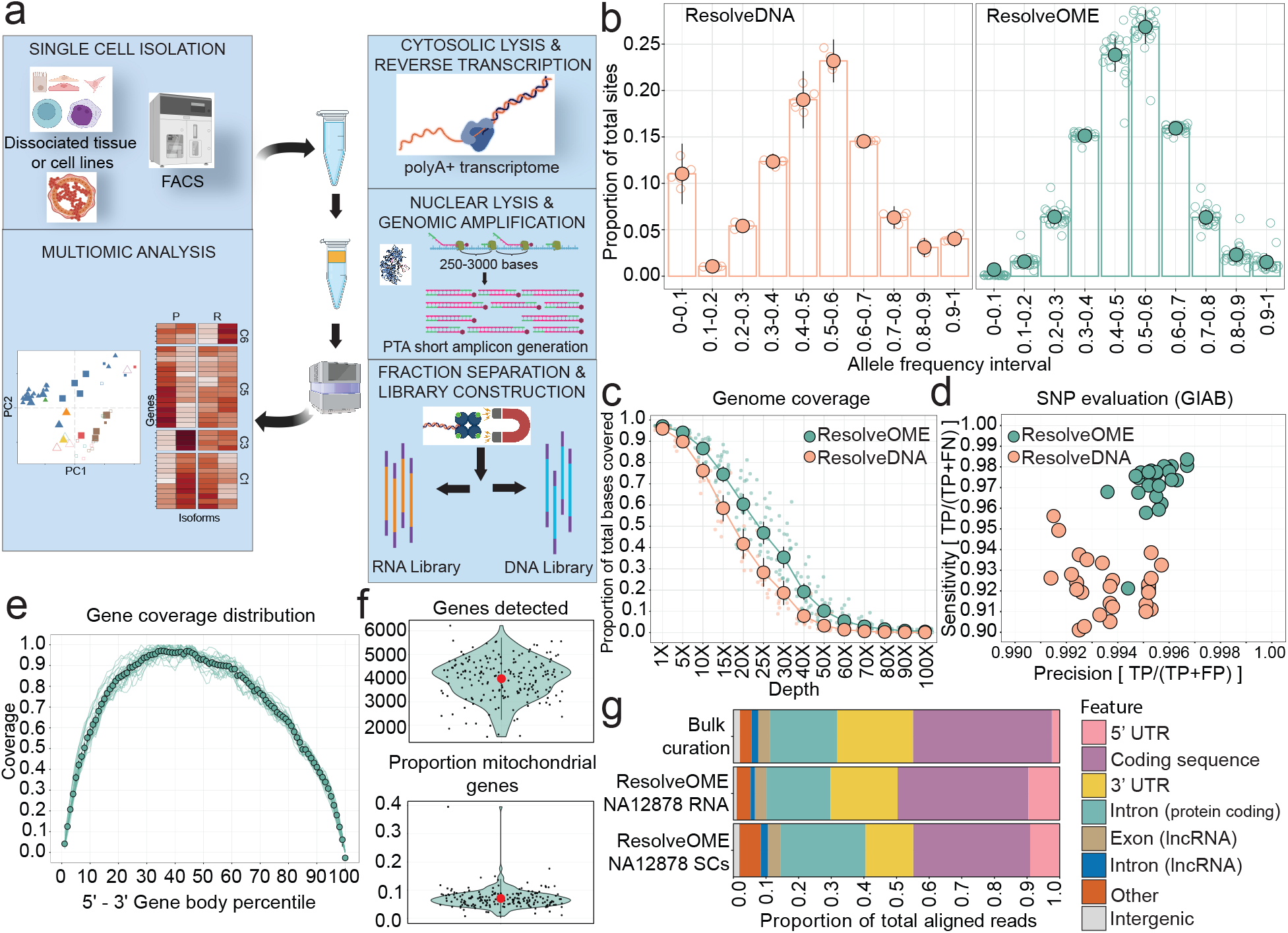
Workflow performance characteristics. **a,** The high-level workflow of enrichment and preparation of simultaneous RNA and DNA from a single cell. **b,** ResolveOME (green) and ResolveDNA (orange) allelic balance in control (NA12878) is shown in deciles of observed allele frequency (AF) across known heterozygous positions. Each dot represents the proportion of variants that showed an AF within the bin frequency for a given cell. Bar plots with error bars describe general trend for all cell-replicates for each AF bin. Allelic dropouts are called when AF is < 0.1 or > 0.9. **c**, Cumulative genomic coverage plot for each sample type performed on ResolveOME, showing the proportion of the entire genome covered (y-axis) at a given depth (x-axis). Each dot represents a cell replicate within a dataset and error plots denote the variability of coverage at a given depth. **d**, SNV calling sensitivity (y-axis) and precision (x-axis), with respect to GIAB NA12878 reference dataset, across the two chemistries (ResolveDNA/PTA, salmon, ResolveOME, turquoise). **e**, Summarized coverage plots for all detected transcripts across the full-length transcript RNAseq chemistry of ResolveOME. X axis is a normalized fraction of a transcript from 5’ to 3’, breaking regions into mean depth per percentile of transcript and y-axis are counts. **f**, Mean number (+/- 2SD) of genes detected from the transcribed nuclear genome (left) or transcribed mitochondrial genome (right) from single cell amplification of NA12878 cells. Single cells are denoted as points; shape of curve indicates overall distribution using the probability density function. **g**, The proportion (averaged across all samples of a group) of aligned reads that matches a specific transcript feature or RNA species is reported for each dataset of single-cell or bulk RNA amplification. Features and proportions were derived from Qualimap summarizations of our transcriptome definition file. NA12878 cells were leveraged, and bulk data was pulled from online repository to serve as reference from typical RNA-Seq. All RNA datasets were down sampled to 100,000 total reads.

We observed that sequencing libraries from DNA showed good complexity using the PreSeq count algorithm^14^. This showed counts of 3.81E9 (+/- 2.95E8) (**Supplementary Figure 1a**) and complexities similar to published reports on NA12878^15^. Cells that showed values greater than 3E9 (90% of all cells) were sequenced at ∼20X genome depth. Confirming good library complexity, allelic balance in the DNA arm showed 98.4% (+/-1.14%), comparable to the 88.2% (+/-4%) for PTA^6^, across 25 replicates (**Figure 1b**). We also observed similar genome coverage distributions between the methods (**Figure 1c**). Using NA12878 benchmarking^16^ reference for GRCh38 we observed improved SNV from PTA^6–8^ with sensitivity of 97.1% (+/- 1.24%) and precision > 99.3% (**Figure 1d**).

In our study, the RNA arm of the chemistry successfully amplified transcripts across the entire gene body of all isoforms detected in NA12878 cells (**Figure 1e, see Methods**). We assessed the performance of RNA quality using established references, including SEQC/MAQC-III Consortium (2014) and Chen, W. et al. (2021), comparing our samples to well-characterized RNA reference standards (Human Brain Reference RNA and Universal Human Reference RNA)^17,18^ (**Supplementary Figure 1b**). We observed similar patterns in NA12878 and reference cells, with mitochondrial read percentages below 10%, indicating effective amplification of polyadenylated transcripts and the inclusion of healthy cells (**Figure 1f, Supplementary Figure 1a**). Furthermore, the workflow also captured various transcript types and components relative to bulk datasets, mRNA-seq alone, and HBRR/UHRR references (**Figure 1g, Supplementary Figure 1b**). Additional quality control metrics at the genomic and transcriptomic levels are provided in **Supplementary Figure 1a**.

### Copy number variation in an acute myeloid leukemia (AML) drug resistance model

MOLM-13 is an AML derived cell line that we treated with the FLT3 inhibitor quizartinib in order to exploit an intrinsic *FLT3* (13q12) mutation to the point of resistance (**Supplementary Figure 2a**). We performed WGS (∼25x) and subsequent CNV analysis on 9 parental “P” and 10 quizartinib-resistant “R” cells. Utilizing a 500 kb window size, copy number gains were evident on chromosomes 5, 6, 8, and 13 concordant with karyotypic data for the parental cells (**Figure 2a**). Highlighting heterogeneity in these untreated P cells, universal 3N ploidy was observed for 9/9 cells for Chr. 5, but only 5/9 cells showed concomitant 5p gains. The R cells exhibited this 5p gain but also surprisingly 7/10 cells showed reversion of Chr. 5p to the diploid state (**Figure 2a**) suggesting this reversion was optimal for or mediating drug resistance. In addition, we observed a novel 19q gain in 4/10 R cells mostly independent of the 6/10 short gains in 6q. Taken together, the R MOLM-13 population exhibited 5p, 6q, and 19q modifications that favored survival under therapeutic pressure.

**Fig. 2.**
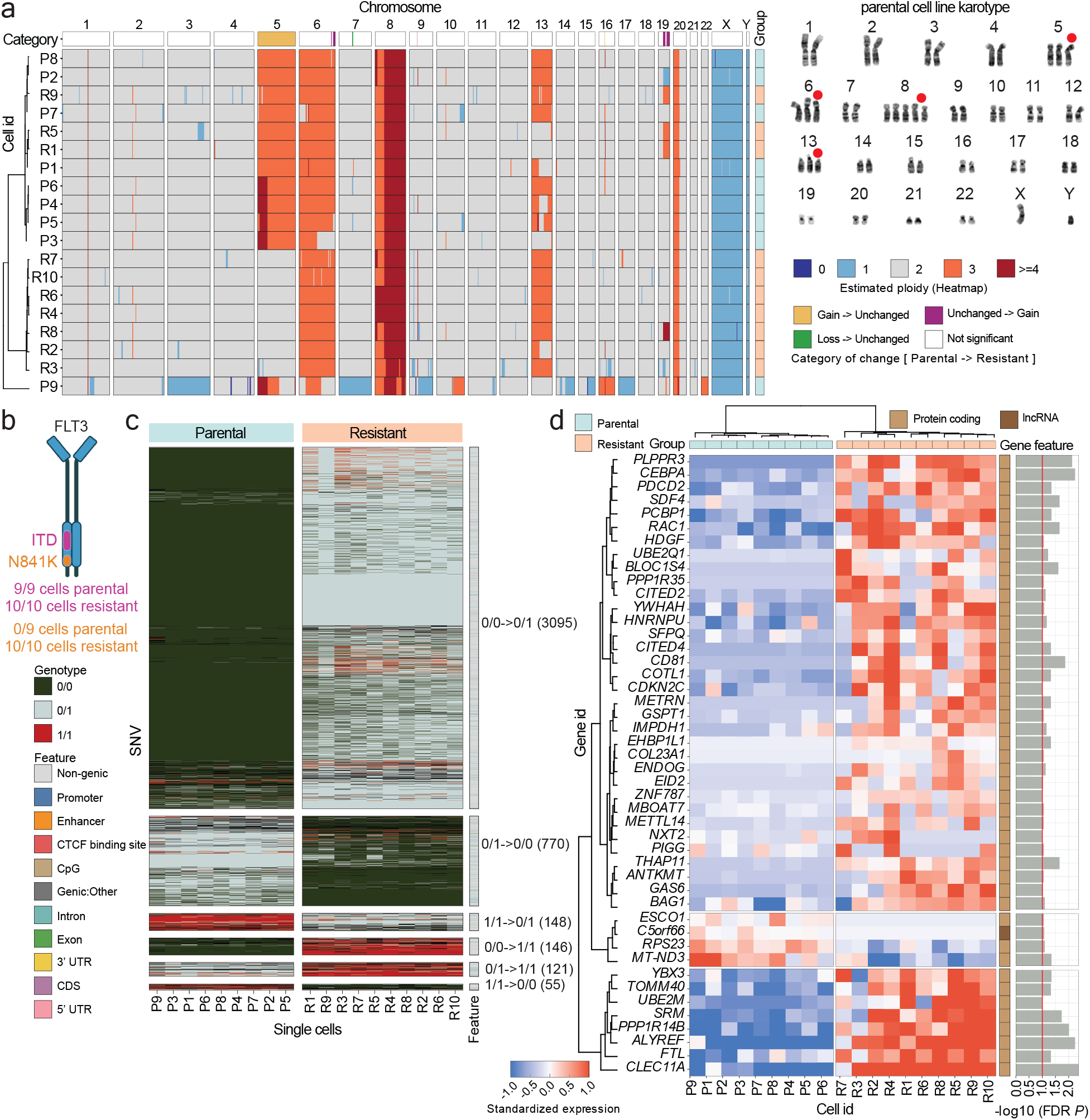
Genomic and transcriptomic alterations in individual parental vs. quizartinib-resistant AML single cells. **a,** Left: Copy number alterations of individual MOLM-13 cells (rows) from parental (turquoise) and resistant (salmon) cells using a bin size of 500kb with Ginkgo. Dendrogram was generated based on distance of each bin’s average fold change from 2N. A summary of the parental to resistant directionality of the copy number alteration (“category”) is shown above each chromosomal segment. Right: Representative metaphase spread of 25 total karyotypic spreads of the parental MOLM-13 line. Red circles denote abnormally amplified chromosomes prior to the acquisition of resistance. **b**, Schematic of FLT3 receptor tyrosine kinase and locations of ITD mutation (juxtamembrane domain) and N841K mutation (tyrosine kinase 2 domain). Prevalence of each mutation between parental and resistant cells is indicated. **c**, Heatmap of SNVs showing statistically significant (p < 0.05 by multinomial logistic regression) genotype prevalence across the MOLM-13 parental and resistant cells. Columns represent cells and rows SNV IDs. Color within the tiles represent the called genotypes. Number of SNVs falling into each category of differential parental/resistant genotype is indicated, as well as ENCODE genotypic feature mapping to the single nucleotide variant (right bar). **d**, Unsupervised clustering of differentially expressed genes in MOLM-13 model of drug resistance. Parental single cells (turquoise) and quizartinib-resistant (salmon) single cells comprise columns; Gene Symbols comprise rows. Gene biotype (protein-coding or lncRNA) and FDR is presented to the right of the heat map; red line indicates q < 0.1.

### Acquisition of a *FLT3* missense mutation as a key driver of drug resistance

We next sought to determine candidate key drivers of quizartinib resistance at the single nucleotide resolution. In addition to the known FLT3 internal tandem duplication (ITD)^19^ in the P cells, an N841K variant was detected in all R cells (**Figure 2b**). This mutation (N841K) in activation loop of FLT3^20^ likely results in constitutive kinase activation^21^ and this mutation and additional activation loop mutations proximal to N841 have expanded during quizartinib treatment in a clinical study^22^. This suggests that this missense mutation is contributing to quizartinib resistance in this model by preventing efficiency of drug binding.

To assess whether the N841K *FLT3* mutation arose *de novo* or was an existing variant cell clone in the parental population we employed a quantitative PCR-based genotyping assay. In parental cells, while amplification of native N841 dominated, a low but detectable level of mutant K841 was detected, compared to resistant cells where the heterozygosity was confirmed through equal C_T_ ratios of N841 and K841 (**Supplementary Figure 2b**). This suggests that the mutant *FLT3* N841K exists as an extremely rare clone in the parental AML cells.

### Heterogenous SNVs in drug resistance

We took the initial approach to review variants previously implicated in AML pathogenesis^12^ with no direct overlap with our quizartinib-resistant cells. Accounting for cellular heterogeneity, we then identified a total of 4335 candidate alleles where genotypes shifted state across P vs. R cells (Wald test, p <0.001) (**Figure 2c, Supplementary Figure 3, Supplementary Table 1**). Of the 4335 candidate alleles with differential genotypic prevalence between P vs R cells, 4003 of them fell in intergenic regions and 332 SNVs within genic boundaries (**Supplementary Table 1**). One notable missense was A109V in the E3 ubiquitin ligase gene RNF167 found in all 10 quizartinib-resistant cells but not present in parental cells.

### Transcriptional signatures of drug resistance

RNA performance metrics were robust for MOLM-13 single cells, exemplified by alignment statistics and exonic and mitochondrial read percentages (**Supplementary Figure 2c**). **Figure 2d** highlights 46 differentially expressed transcripts between the P and R single cells including marked upregulation of *GAS6*, a ligand for the receptor tyrosine kinase AXL (**Supplementary Table 2**). The AXL pathway, specifically through downstream STAT3 cell proliferation and PI3K/AKT survival signaling, has been shown to be a bypass pathway for FLT3 inhibition^19,23^ (Supplementary Figure 2d). We also observed concurrent transcriptional upregulation of the small GTPase *RAC1*, which may be synergistic with upregulation of the AXL-STAT3 and AXL-PI3K/AKT signaling axes^24,25^. Relatedly, we also noted increased expression of the pioneer transcription factor CEBPA CCAAT/enhancer-binding protein alpha (C/EBPα) in quizartinib-resistant cells. Truncating mutations in *CEBPA* are found in ∼10-15% of AML patients^26,27^ leading to expression of an N terminal fragment of CEBPA, p30 with potential dominant negative^28^ activity.

### Isoform switching in drug resistance

Leveraging the 5’:3’ transcriptome coverage of our approach, we performed differential transcript usage (DTU) analysis (**Figure 3a**) to identify alternative splicing at the single cell level. We identified 55 genes for which two major isoforms were differentially utilized in the resistant cell cohort including *CASP3* isoform “B” that was selected for in the resistant cells relative to parental cells, potentially contributing to alteration of apoptotic programs and contributing to drug resistance. **Figure 3a** highlights an additional 3 genes suggestive of functional impact in AML: cell hematopoiesis (*CD164*^29,30^), DEK-mediated oncogene (*BANF1*^31^) and chemotherapy resistance (*VDAC2*^32^). Transcript IDs for these 55 genes are in **Supplementary Table 3**.

**Fig. 3.**
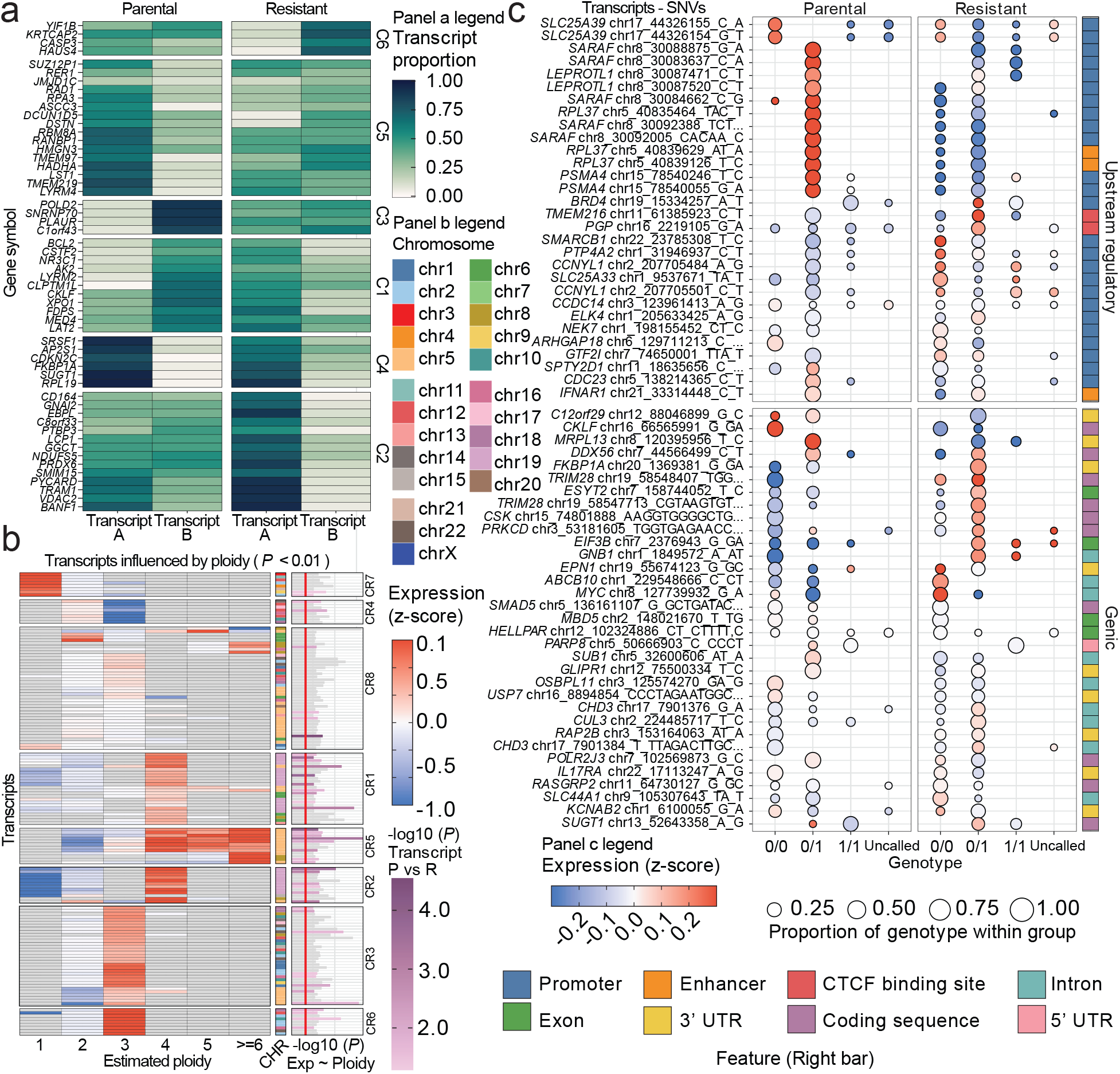
Unification of genomic lesions and gene expression in AML model of drug resistance. **a,** Differential transcript utilization (DTU) between MOLM-13 parental and drug-resistant single cells. Color intensity indicates transcript proportion of A or B isoform of indicated transcript. **b,** Heatmap with transcripts in the y-axis that show a statistical (ZLM p < 0.01) association with ploidy level across all cells in the MOLM-13 dataset. Color of the tiles represents the average standardized expression value at a given ploidy level. The right panel shows the output of the ZLM model testing the expression given the ploidy. Red line indicates the p < 0.05 cutoff of the model. Bars are colored based on the -log10 p-value of the ZLM model testing transcriptional differences between parental and resistant cells. Blocks of concordance (ploidy and expression increased or decreased concomitantly) or discordance (ploidy and expression inversely correlated) are shown for a given transcript and chromosomal location. **c**, Bubble plot showing SNV-transcript expression associations (p < 0.05) determined by ZLM modeling between parental and resistant cells. Candidate SNVs are shown in the y-axis and genotypes in the x-axis. Size of the circle denotes the genotype prevalence of the variant in the MOLM-13 cell type set (parental or resistant). Colors of points denote the standardized mean expression level of the transcript in the set. ENCODE genotypic features mapping to the given single nucleotide variant are indicated in the right bar and are categorized in the heatmap as regulatory (top) or genic (bottom).

### Relationship of gene expression to copy number

Concomitant with the Chr 19q gain, we noted transcriptional upregulation of *CEBPA* (19q13.11) in a subset of R cells (**Figure 2a**). Other cells demonstrated increased transcript levels without copy number gains at this locus suggesting other mechanisms leading to overexpression. In support of this, we identified a candidate distal promoter/enhancer SNP ∼20kb 5’ of the *CEBPA* transcriptional start site with a genotypic bias between parental and resistant cells (**Supplementary Table 4**).

These instances prompted us to examine the relationship of ploidy to gene expression across the genome. We identified a set of 341 transcripts with statistically meaningful associations of ploidy to expression (p<0.05) (**Figure 3b, Supplementary Table 5**). Intriguingly, this analysis also exposed abundant discordant or inverse gene copy to expression relationships. We found clusters of genes that were reduced in copy number yet were overexpressed in R vs. P cells (**Figure 3b**, CR5), and, conversely, clusters existed where genes were under expressed in instances of ploidy gain (**Figure 3b**, CR4, CR7 and CR5). This cautions against copy number inference based solely on gene expression data, conversely, not all copy number variations result in coordinate changes in gene expression.

### Identification of candidate regulatory SNVs modulating transcript levels in drug resistance

As we identified occurrences of genomic lesions of interest that did not associate with the predicted transcriptional levels, we sought to identify proximal single nucleotide variants (-5000 to 0 relative to the transcriptional start site, or intragenic) associated with differential expression of the linked gene in P vs. R cells (**Figure 3c**, **Supplementary Table 6**). We highlight 63 regulatory alleles where both gene expression and genotype are significantly different between parental and resistant cell cohorts. These single nucleotide variants resided in different regulatory elements, including promoters, enhancers and CTCF binding sites. A putative promoter variant in the *BRD4* gene, a master transcriptional regulator, was identified as a candidate SNV influencing increased *BRD4* expression in resistant vs parental cells. Intriguingly, we identified an intronic variant in *MYC* which was associated with upregulation of *MYC* expression in resistant cells. Regulatory variant heterogeneity between and within parental and resistant single cells is only exposed with unified genomic and transcriptomic profiles from the same individual cell.

### Primary DCIS/IDC single cells exhibit heterogeneous classes of chromosomal loss

We next applied our methods to disassociated cells derived from a primary human breast cancer containing both ductal carcinoma in situ (DCIS) and invasive components (estrogen and progesterone receptor positive, HER2 negative). Normal breast cells from this same individual were analyzed for comparative purposes. Single cells were enriched for tumor cells by FACS sorting based on expression of the epithelial specific cell surface marker, EpCAM.

Distinct classes of CNV emerged, where single cells exhibited discrete chromosomal losses. We found a subset of single tumor cells (16/22) that harbored a 13q loss with concurrent loss of 16q/17p (**Figure 4a**). The most abundant class (11/22 cells) harbored these copy number alterations plus a third discrete loss of Chr. 11q. The observed 13q and 16q/17p loss is consistent with reported copy number alteration in multiple stages of breast cancer advancement^33^ and coincides with the loss of the prototypical tumor suppressor genes *BRCA2, RB1* and *TP53*. Interestingly, we observed gain of Chr. 13p, a heterochromatic “stalk” devoid of genes in 9/22 tumor cells, and Chr. X gain of unknown significance in 5 cells total encompassing the centromere and flanking p and q arm segments. Even with this small cohort of single cells, these data are representative of a heterogeneous tumor undergoing active evolution.

**Fig. 4.**
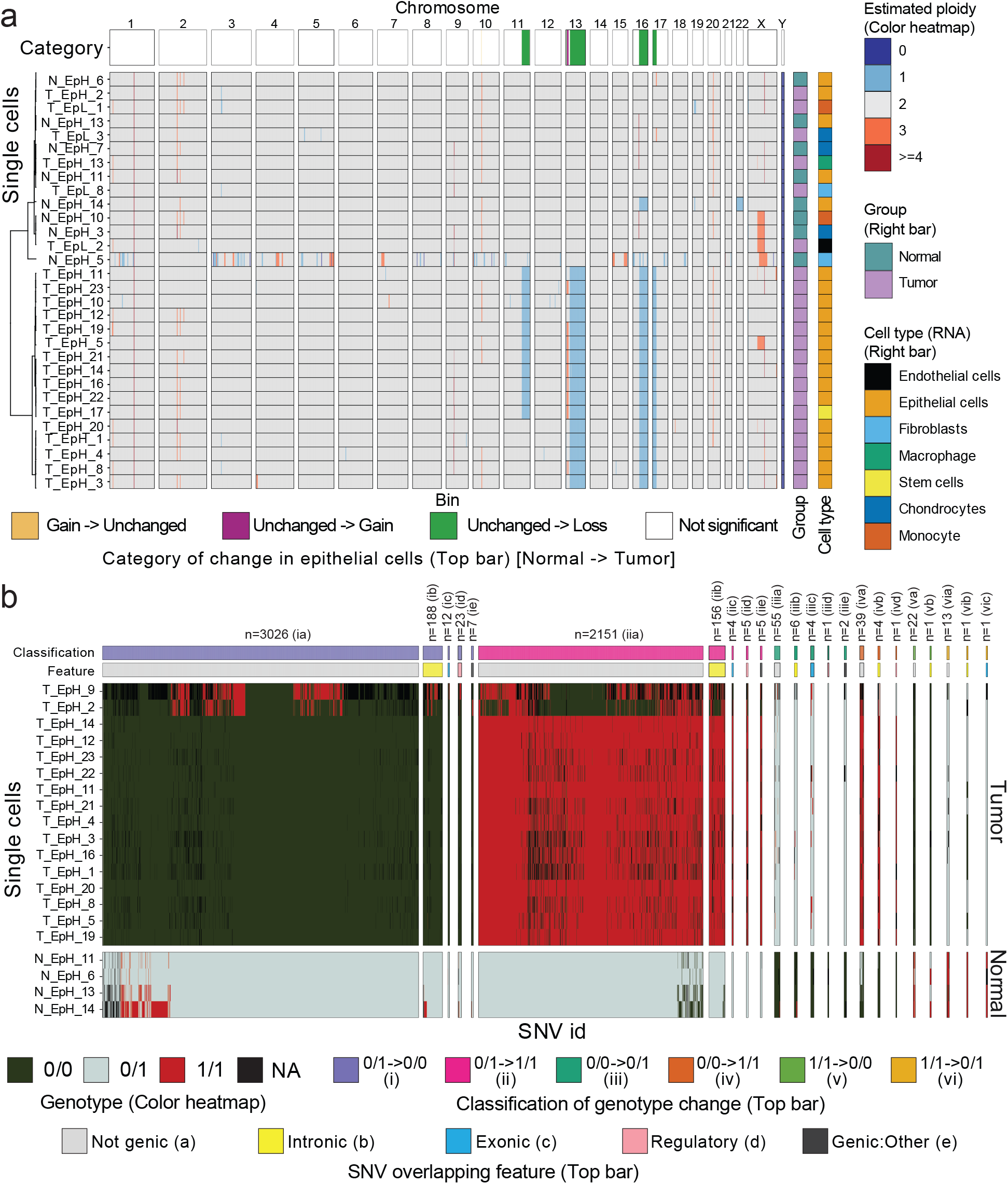
Copy number alterations and single nucleotide variation in patient breast cancer single cells. **a**, Copy number alterations in primary DCIS/IDC from tumor biopsy (purple) or matched normal biopsy (green) single cells. Status of EpCAM FACS (High or Low) categories is indicated in the sample name. Two distinct classes of chromosomal loss are observed in EpCAM high cells from the tumor fraction: 1) combined 11q, 13q, 16q/17p loss and 2) combined 13q and 16q/17p loss. Additionally, 13p gain was identified in 9/22 tumor cells, while Chr. X gain encompassing the centromere and flanking P & Q segments was noted in 5 single cells. Cellular identity called by transcriptional profile is indicated (right bar) **b**, Differential genotypic prevalence in tumor vs matched normal biopsy cells upon deep sequencing. The heatmap displays single cells (rows), single nucleotide variant (columns), and corresponding 0/0 homozygous reference, 0/1 heterozygous, or 1/1 homozygous alternate genotype status (color). Discrete SNV blocks of the heatmap (left and right) are observed for 0/1 heterozygous identity for normal biopsy cells with corresponding homozygous identity for tumor cells, influenced by copy number loss in the tumor cells. The direction of the genotypic change is colored by class and SNVs are colored by genomic feature (top bars).

### Single nucleotide variation in DCIS/IDC

Prior to genome-wide unbiased assessment of SNV, we assessed exons of the *PIK3CA* gene, one of the most frequently mutated genes across diverse molecular subtypes of breast cancer. We identified the missense mutation N345K in 14/22 cells from the tumor biopsy, all of which were classified by FACS as EpCAM high but did not identify this variant allele in the EpCAM low cohort or in cells from the normal breast tissue. The N345K mutation is second only to H1047R amongst *PIK3CA* hotspot mutations catalogued by TCGA^34^ and is known to influence the interaction of the p85 (*PIK3R1*) regulatory/p110 (*PIK3CA*) catalytic subunits by disruption of the C2/iSH2 domain interface^35–37^.

Surveying the coding sequence of genes known to be influential in ER+ breast cancer^11^, we identified a Q73H missense mutation in the transcriptional cyclin *CCNT1* (15/17 EpCAM high tumor cells, 0/4 EpCAM high normal cells) and a missense mutation in *CDH2* R753S (16/17 EpCAM high tumor cells, 0/4 EpCAM high normal cells). Genetic variants detected in tumor EpCAM high vs. ipsilateral normal EpCAM high cells are summarized in **Figure 4b** and **Supplementary Table 7**.

### Cell identity and transcriptional state of DCIS/IDC singulated cells

Differential gene expression analysis highlighted gene signature blocks between a primary clade of exclusively EpCAM high cells from the tumor biopsy, and clusters comprised of intermixed EpCAM low/high cells and intermixed tumor/normal status (**Figure 5, Supplementary Table 8**). Leveraging cell type identification (**see Methods**) we defined the cell type diversity. Intriguingly, an EpCAM high tumor cell that harbored *PIK3CA* N345K and copy number alteration profiled as a stem cell and exhibited hallmarks of an EMT gene expression signature (**Figure 6**), including differential expression of *SPINT2*^38^*, CDH1*^39^*, and ERBB3*^40^ relative to the other epithelial cells. Additionally, one outlier EpCAM high tumor cell possessed oncogenic *PIK3C*A, an under expression of *VIM,* and the prototypical breast cancer chromosomal losses yet profiled closer to normal biopsy cells by gene expression signature, also suggestive of a potential epithelial-to-mesenchymal transition.

**Fig. 5.**
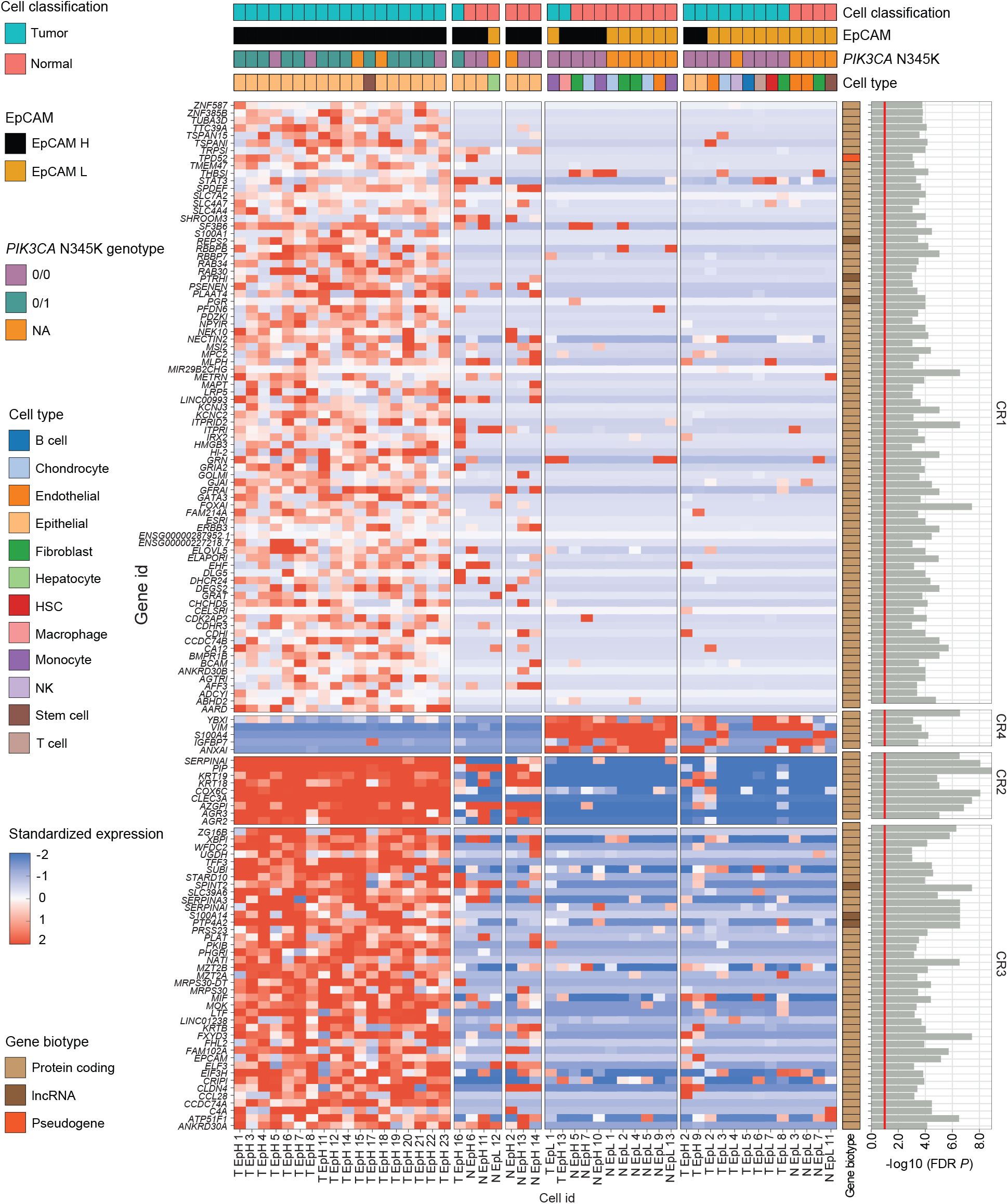
Multi-omic integration of individual primary breast cancer cells. Unsupervised clustering yields four primary signatures (CR1-CR4) of differential gene expression from a single cell sample set derived from tumor or matched normal biopsy specimens. EpCAM FACS status (high or low), *PIK3CA* (N345 wildtype or K345 mutant) genotypic status, cellular identity call, and gene biotype (protein-coding, lncRNA, or pseudogene) are shown for each single cell (column). P value significance (right bars) is presented for each transcript (row). EpCAM High tumor cells predominantly type epithelial by gene expression and harbor *PIK3CA* N345K, while EpCAM Low cells from both tumor and matched normal biopsies represent a diversity of cell types with wildtype *PIK3CA*.

**Fig. 6.**
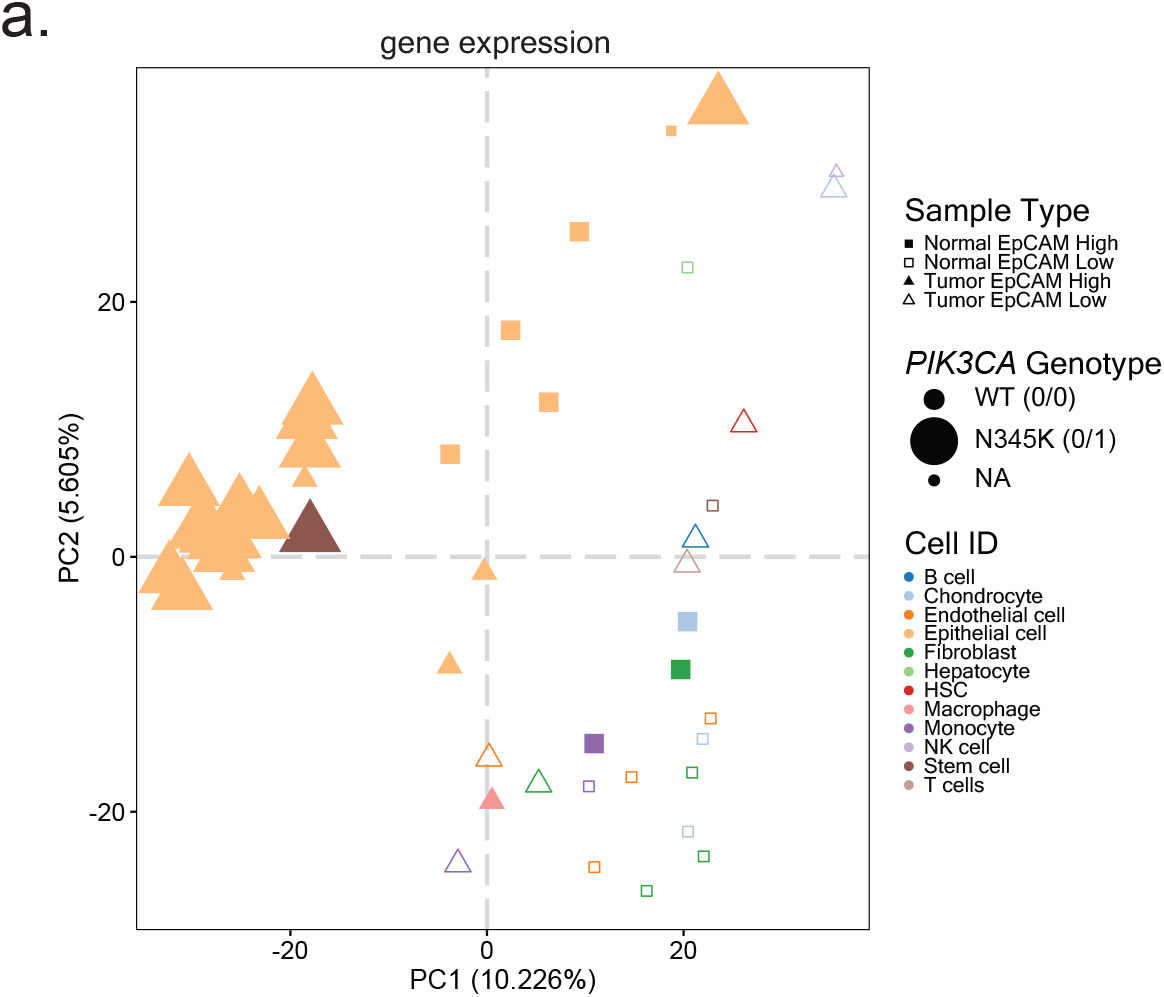
Single cell identity and state plasticity in the context of *PIK3CA* genotype. Principal component analysis of single cell datasets separates tumor (triangles) and matched normal biopsy (squares) cells by gene expression. EpCAM High cells (solid icons) enriched by FACS type predominantly epithelial (yellow) while EpCAM Low (open icons) cells highlight the extensive cell type diversity (colors) present in the biopsies. Most tumor epithelial cells harbor oncogenic *PIK3CA* N345K (large icons) while normal control biopsy cells are wildtype (medium icons), and these classes are expectedly separated in PCA space by gene expression. One outlier cell (large brown solid triangle) harboring a *PIK3CA* N345K mutation types stem-like vs epithelial, indicative of an intermediary cell state, potentially EMT.

## DISCUSSION

Cancer is a genetic disease driven by alterations that are translated through the transcriptome to the proteome manifesting as tissues with aberrant function. These diverse changes, some of which have been characterized, result in remarkable heterogeneity amongst the cells that populate primary and metastatic tumors. Development of a robust single-cell multi-omic analysis allows for comprehensive concurrent examination of tumor heterogeneity, which is required to define the evolution and pathogenic trajectory of the disease. First, we have demonstrated robust allelic representation underlying cellular identity along with profiling the transcriptome. Second, we exemplify the plasticity of the genome and consequential transcriptome state when tumor cells are subjected to a selective pressure. Third, we demonstrate the unbiased identification of malignant cellular clones with synced multiple molecular modalities which can increase sensitivity of molecular lesion detection.

The generated data using the NA12878 cell line enabled us to determine the reproductivity and accuracy of the transcriptome analysis as well as assess the performance of the genome amplification arm of the single-cell multi-omic chemistry. We found the gene body detection was improved when compared to 3’ end counting single-cell RNAseq methodologies and demonstrated impactful isoform representation. Moreover, we found the performance of both CNV and SNV detection was either equivalent or actually improved over the previously developed single-omic genome-based analysis^6^ which is driven by the high allelic balance and low allelic drop-out. This high coverage and strong allelic balance culminated in the ability to call SNVs with confidence. Having demonstrated the accuracy, sensitivity, and precision of the method we then focused upon our ability to delineate the molecular basis of drug resistance in using a patient derived cell line model. We were struck by the single nucleotide variation present in our parental vs. quizartinib resistant MOLM-13 cells (4335 differentially prevalent SNVs, **Figure 2c**, **Supplementary Table 1**), which further underscores that, while transcriptional plasticity is dogmatic, it is equally as important to recognize *genome* plasticity. Efforts to estimate intergenic variation at putative functional elements—promoters, enhancers, splicing enhancers—is a frontier and an underappreciated aspect of drug resistance studies. In single cell tumor biology, most work to date has been performed at the transcriptome level, owing to the large-scale adoption of droplet-based methodology facilitating workflow ease and single-cell throughput. While there has been unquestionable advance from droplet-based RNA-Seq studies defining diversity and heterogeneity of cell types, a gap remains in that there have been few studies providing genetic determinants of these cells’ programming. In the absence of DNA-level information, mechanisms of transcriptional impact, such as variation in regulatory elements, in transcription factor binding sites, or in chromosomal copy number remain undefined. Secondly, while transcript-level information is frequently employed for molecular subtyping of a tumor^41,42^, pharmacological and diagnostic decisions are primarily driven by genomic variation due to technical and informatics challenges with ascertainment by transcriptional status^43^.

Joint RNA/DNA single-cell profiling has enabled us here to spotlight instances of diverse cell types in our primary breast cancer sample. This workflow negates the need to enrich or bias a heterogenous tumor specimen with a biomarker—the genotypic diversity of all cells in the sample can be ascertained and considered together as microenvironment interactions as the gene expression profile provides the cell typing to interpret genotype.

Analysis of single cell data derived from primary human tumor and matched normal specimens revealed previously unrecognized copy number and single nucleotide variations which accompany and may have driven the evolution of this tumor from pre-invasive DCIS to invasive cancer. In addition, an intermediate transcriptional profile was intriguingly observed in an EpCAM high cell that harbored *PIK3CA* N345K and DCIS/IDC-characteristic chromosomal losses^33^, thus having the core genomic changes of the main epithelial cell cohort. Nevertheless, it manifested with a different transcriptional stem-like state with hallmarks of EMT signatures—indicating a potential state conversion^44^ as well as highlighting inherent transcriptional single-cell heterogeneity even within a relatively small sampling of a singulated tumor sample. It will be crucial to define the diversity of additional novel transcriptional states that may be contributing to the advancement of DCIS to invasive cancer.

Each “-omic” tier of molecular information allows a greater ability to comprehensively define the molecular mechanisms of oncogenesis in a tumor. The ability to simultaneously analyze genomic and transcriptomic data from the same individual cell vastly increases the complexity of putative mechanisms of drug resistance and oncogenesis and represents a vast computational biology challenge. This will only increase as additional “-omic” tiers of layers are added, including ascertainment of protein expression as the nature as workflows evolve to allow for the incorporation of CITE-seq-like^45^ oligo-tagged antibodies. We envision mechanistic insights analogous to those presented here to accumulate from the research community having the newfound capability to accurately assess single nucleotide genomic variation in conjunction with transcriptional profiles—aiding discovery efforts to generate a new wealth and generation of pharmacological targets.

## METHODS

### Cell culture

NA12878 cells (CEPH/Utah Pedigree 1463) were obtained from the Coriell Institute for Medical Research (Camden, NJ). Cells were maintained in RPMI 1640 (Gibco 11875-093) supplemented with 15% FBS and penicillin/streptomycin, and sub-cultured every 2-3 days while maintaining a density range of 2.5 E5 – 1.5 E6 cells/ml.

MOLM-13 acute myeloid leukemia cells harboring heterozygous FLT3 internal tandem duplication (ITD) were obtained from the DSMZ-German Collection of Microorganisms and Cell Cultures (ACC 554). Cells were maintained in RPMI 1640 (Gibco 11875-093) supplemented with 10% FBS and penicillin/streptomycin, and sub-cultured every 2-3 days while maintaining a density range of 2.5 E5 – 1.5 E6 cells/ml. For generation of the quizartinib-resistant MOLM-13 line, cells were continually treated with 2 nM quizartinib (SelleckChem AC220) or DMSO vehicle control for matched parental control line and drug replenished at each subculturing until emergence of resistant clones at 5 weeks duration in culture. Genomic DNA (Zymo Research Quick-DNA Microprep Plus Kit, D4074) or total RNA (Qiagen RNeasy Plus Kit, 74034) was isolated from quizartinib-resistant and matched parental MOLM-13 cells at time of FACS sorting to generate bulk sequencing control libraries for comparison to single cell datasets and for quantitative PCR template.

### Multi-omic amplification workflow

Singulated cells were processed using the manufacturer’s specifications for ResolveOME. This begins with template-switching-based RNA-Seq chemistry to generate biotin-dT-primed, first strand cDNA followed by termination of the reaction and nuclear lysis, at which point primary template-directed amplification proceeds. The mRNA-derived cDNA is affinity purified with streptavidin beads from the combined pool of cDNA and amplified genome. cDNAs are then further purified with subsequent streptavidin bead washes of two stringencies and on-bead pre-amplification of the first-strand cDNA to yield double-stranded cDNA. In parallel, the PTA fraction from the same cell containing genome amplification products, separated from the cDNA, is purified. The separate and distinct fractions of pre-amplified mRNA cDNA and genome-derived DNA amplification fractions undergo SPRI cleanup prior to NGS library generation.

### Karyotyping

MOLM-13 cells were analyzed within 2 weeks of thaw (KaryoLogic, Inc, Durham, NC) with a workflow for complex hyper diploid karyotypes using 25 metaphase spreads. Live cultures were delivered to the service provider (KaryoLogic, Durham, NC) on-site and cultures recovered in 5% CO2 37C incubators on-site for one week prior to metaphase spread creation.

### FACS

Prior to FACS, cell lines were first counted and assessed for overall viability by trypan blue staining using a Countess II FL instrument (ThermoFisher Scientific) or by acridine orange + propidium iodide with a Luna FL instrument (Logos Biosystems). Cell line cultures put forth to the FACS protocol exhibited >90% viability.

### MOLM-13

For single cell analysis, ∼2.0E6 MOLM-13 quizartinib-resistant or matched parental cells were rinsed twice in staining buffer (0.2 µm filtered Dulbecco’s Phosphate Buffered Saline lacking calcium and magnesium (Gibco 14190) supplemented with 2% FBS) and kept on ice until BD FACSAria III sorting at the UNC School of Medicine Flow Cytometry Core Facility. Following Calcein AM (BioLegend 425201), propidium iodide (Millipore Sigma P4864) and DAPI staining, singlet (FSC-A / FSH-H, SSC-A / SSC-W) and live cell (DAPI/PI negative, top 70% Calcein-AM positive) gating was established and single cells were sorted (130 micron nozzle assembly) into low-bind 96 well PCR plates (Eppendorf twin.tec LoBind, semi-skirted, 0030129504) containing Cell Buffer from the ResolveOME kit and immediately frozen on dry ice following centrifugation.

### NA12878

∼2.5E6 NA12878 (NA12878/HG001) cells were prepared as above and subjected to Sony SH800 sorting using a 130-micron chip. Singlet (FSC-A / FSC-H, BSC-A / BSC-W) and live-cell (PI negative, top 70% Calcein-AM positive) gating was employed for single cell sorting into low-bind 96 well PCR plates pre-loaded with Cell Buffer from the ResolveOME kit as described above.

### Primary DCIS/IDC

Tissue for single-cell DCIS/IDC studies was obtained in accordance with the Duke University Medical Center IRB for the clinical trial PRO00034242 “Biologic Characterization of the Breast Cancer Tumor Microenvironment.” Cryo-preserved, singulated cells (∼4.2E5) derived from tumor or matched normal mastectomy tissue were thawed at 37C and centrifuged at 350 x g for 5 min to separate cryo-preservation media. Cells were rinsed once in staining buffer and incubated with 2 µg/ml anti-human CD326 conjugated with AlexaFluor 700 (ThermoFisher 56-9326-42) at 4C in the dark for 1h. Following this, ∼8.4E4 cells were reserved for a parallel negative control mock stain lacking any antibody for assessment of background fluorescence levels for viability and EpCAM staining. Then cells were washed 3X with staining buffer with 350 x g 5 min centrifugations in between washes and passed through a 35-micron filter prior to loading for FACS. Singlet (FSC-A / FSC-H, BSC-A / BSC-W) and live-cell (Calcein AM, BioLegend 425201) gating was defined followed by daughter EpCAM high and EpCAM low gates. EpCAM High and Low cells were sorted into the same 96 well plates as described above to minimize potential batch effects of downstream genomic/transcriptomic amplification.

### Quantitative RT-PCR

10 ng of genomic DNA was isolated from a cell collection of quizartinib-resistant or matched parental cells as described above and subjected to a custom Taqman™ genotyping assay, #ANMF9C4 (Invitrogen-Applied Biosystems) using the manufacturer’s suggested conditions for reaction assembly and cycling on a QuantStudio6 instrument. The assay was designed to distinguish between human N841 and K841 with the C/A nucleotide polymorphism, respectively at the GRCh38 / hg38 coordinate Chr13:28,018,485.

### Sequencing

Low-pass sequencing was first performed on the DNA arm libraries using an Illumina MiniSeq (2.3 pM library flow cell loading concentration) or NextSeq1000 (640pM library flow cell loading concentration), 2X75 targeting >2.0E6 total reads per library. For RNA fraction libraries, 2X75 MiniSeq or NextSeq1000 sequencing targeting on average >1.0E6 reads per library was employed for flexibility for data down-sampling. For joint clustering of DNA and RNA fraction libraries, a 4:1 molar ratio of [DNA arm]:[RNA arm] libraries was employed. Following low-pass sequencing, the DNA arm libraries were 2X150 sequenced on an Illumina NovaSeq6000 S4 flow cell targeting 5.5 E8 total reads to provide down-sampling flexibility at either the Vanderbilt Technologies for Advanced Genomics (VANTAGE) core facility or the Duke University Genomics and Computational Biology (GCB) core facility.

### Bioinformatics Approaches

#### Benchmarking multi-omic data quality evaluations

##### Pre-Sequencing DNA quality control

To evaluate the single-cell libraries’ suitability for high-throughput sequencing, BaseJumper^TM^ DNA-QC pipeline v1.7.3, which uses low-pass sequencing data to construct numerous quality control measures, was used. Particularly, we obtained the pre-sequencing read count to gauge library complexity. This pipeline also provides measures for genomic coverage, the percentage of reads that map to chimeras, the percentage of reads that map to mitochondrial DNA (chrM), the percentage of reads that align to the reference genome, and the percentage of nucleotides that are mismatched to the reference genome. Additionally, the pipeline leverages MultiQC for supplemental QC metrics such as read length, % of duplicate reads, number of mapped reads, and total number of mapped reads.

##### Evaluating RNA fraction quality

Following low pass sequencing, quality control was performed using HG001 NA12878 B-lymphocyte cells to benchmark estimations of quality for the RNA fraction of the workflow. Several quality measures were considered for evaluating data quality, repeatability, and overall protocol robustness using the BaseJumper^TM^ BJ-Expression pipeline v1.6.1. For each cell the percent of reads mapped to the reference genome, as well as the percentage of read aligned to exonic and intergenic genomic features were quantified leveraging Qualimap^46^ (v2.2.2) platform as an internal module of the BJ-Expression pipeline, for bias estimations as proportion of total filtered input reads. Likewise, characteristics like the number of protein coding genes/transcripts identified, dynamic range, and transcript body coverage were utilized to assess assay performance, robustness, and uniformity of coverage. The number of detected genes is defined as the number of protein-coding genes with at least one expression count as estimated by the Salmon^47^ (v1.6.0). Full-length reads, particularly in protein-coding areas, are termed detectable if at least one full-length read count is read by the completely mapped gene. The dynamic range refers to the span between the highest and lowest expression levels of genes across all individual cells in a dataset. It represents the magnitude of variation in gene expression across the cellular population^17^. The top and bottom 10% of gene expression quartiles across normalized counts per million for each cell’s gene expression levels are for this estimate. The uniformity of transcript read coverage in the RNA fraction is defined as the number of reads that cover each nucleotide position along the gene transcript body’s 5’ to 3’ length.

### High Pass DNA Sequencing Evaluations

#### Variant detection evaluation

The BJ-WGS BaseJumper^TM^ pipeline v1.6.0, which comprises of the Sentieon GVCFTyper, VarCal, and ApplyVarCal modules, was used to perform joint genotyping for NA12878 cells. The sensitivity and precision of called SNP variants were then determined using the variant calling evaluation module of the BJ-WGS pipeline, which provides variant quality score log-odds (VQSLOD), with the NA12878/HG001 genome v3.3.2. from the Genome in a bottle (GIAB) consortium as a reference^13^.

#### Allelic balance in NA12878 cells

To appropriately represent allelic balance for each heterozygous site, we determined the ratio of read counts supporting the reference and alternative alleles, using BaseJumper ADO sub workflow (rev. a7240e8) based on a series of bcftools commands that extract the a priori defined GIAB NA12878/HG001 NIST reported high-confidence heterozygous sites. The variant allele depth was retrieved and transformed into a proportion for each cell and heterozygous site. For final reporting, heterozygous sites with a total depth greater than one were selected.

### Secondary multi-omic data analysis

For analyses coming from the DNA fraction of the workflow, we leveraged BaseJumper^TM^ BJ-WGS analytics pipeline (v1.6.0) modified from Sentieon driver-based tools. Initial FASTQ pairs were trimmed against low quality and library artifacts using fastp^48^ (v0.20.1). Alignment was performed using BWA (Sentieon-202112), followed by deduplication (locus_collector v202112 / dedup v202112) of identically aligned reads. Alignment-based QC and coverage determination was (driver_metrics v202010). Copy number calling was performed using ginkgo^49^ (GitHub commit: 892b2e9f851f71a491cade6297f74f09f17acf4c), with a window size of 500kb. Variant calling at the cell level was performed with haplotyper (v202010). Characteristics for all variants were provided for variant quality score recalibration to VARcall, GVCFtyper (v202010). All variant identification and annotations for gene/coding effect were performed using snpEFF/SnpSIFT^50^ (5.0e). Further variant-based tertiary analysis used filtered genomic loci with sequencing depths >4 and >1 variant read candidate SNVs. All candidate SNVs were classified according to allele frequencies.

The BJ-Expression RNA-Seq pipeline (v1.6.1) implemented here was used to generate metrics of feature quantification at the transcript and gene-level. Details about the number and length of reads generated is found in Supplementary Table 1. Unless specified to be down-sampled (using seqtk^51^ v1.3), all reads were leveraged for each analysis. To remove low quality sections and sequencing artifacts, fastp was used for all cells’ analysis prior to alignment. Alignment of reads was performed with STAR^52^ (v 2.7.6a) and were compared against transcript reference made from combining Ensembl (release 104) known transcripts and noncoding. Region assignment and counting of aligned reads was performed with HTSeq4949 (v 0.13.5) and Salmon5050 (v1.6.0) for gene-level metrics. Further, we used the pseudo-alignment algorithm implemented in Salmon to perform both transcript-level and gene-level quantification. Matrices of feature expression were constructed using the Bioconductor package tximport.

### Tertiary multi-omic data analysis

#### RNA fraction: matrix normalization

The logarithmic norm approach was used to normalize the matching Salmon-based transcript and gene matrices for both MOLM-13 and DCIS cells. In summary, feature counts for each cell are divided by total counts for that cell, multiplied by the scale factor (10^4^) and then log2 converted. These normalized matrices were used as input for downstream analyses such as principal component analysis (PCA), differential transcript expression (DTE), differential gene expression (DGE), differential transcript usage (DTU), heatmap reconstruction with unsupervised cell and transcript/gene clustering, and zero inflated linear models linking transcript expression to CNV and SNVs.

#### Principal component analysis

We used the R function scale to center MOLM-13 and DCIS normalized transcript level and gene level matrices across samples within a feature. Principal component analysis was then performed using the oh.pca function from the ohchibi R package, with the centered normalized matrices as input. The PCA model was performed with weighted sparsity constraints due to the heterogeneity of sparsity imposed by the number of transcripts/genes for DCIS normal and tumor cells.

#### Differential expression

We estimated differential transcript and gene expression using the zero-inflated linear model (ZLM) implemented in the MAST^53^ R package using the log normalized feature matrices mentioned above as input. To identify transcripts/genes with signatures of differential expression across parental and resistant cells in the MOLM-13 dataset, we fitted the following model: Transcript/Gene expression ∼ Cell Type (Parental/Resistant) + Number of detected features (transcripts/genes) per cell.

The top 500 highly variable genes in the DCIS dataset were used for principal component analysis, which was the initial step in the process. Non-negative and sparse cumulative PCA was utilized to identify dissimilarities between tumor type (normal and tumor) and EPCAM sort (High/Low) in order to account for the sparsity in the number of genes per cell that exists between tumor and normal. To find transcripts and genes with signatures of differential expression across the aforementioned groups, PC discretized groups were fitted the following ZLM: Cell Group Transcript/Gene Expression.

#### Cellular typing

Transcriptome-based cellular typing was performed for the DCIS dataset using the BJ-Expression pipeline (v1.6.1) cell typing module, utilizing the Human Primary Cell Atlas expression reference dataset deposited in the celldex^54^ R package and cell identification using Seurat^55,56^ R package, by taking as input the gene level normalized expression salmon-based matrix.

#### Differential transcript usage

For the MOLM-13 dataset, we performed differential transcript usage as previously described^57^. Briefly, we took the scaled TPM metric output from tximport and reconstructed a matrix of transcript abundances across cells. Next, we modeled the transcript expression using the Dirichlet-multinomial distribution model implemented in the DRIMSeq R package^58^.

#### Linking transcript expression to CNV

For the MOLM-13 dataset, we linked transcript-level variation in expression with changes in locus ploidy utilizing a zero-inflated linear model framework. Briefly, for each quantified transcript, we extracted its ploidy across cells from the Ginkgo-based estimation by employing genomic-coordinate intersection utilizing the GenomicRanges R package^59^. Next, we fitted the following ZLM design utilizing the MAST R package: Transcript expression ∼ Estimated ploidy at a given locus.

#### Linking transcript expression to genomic polymorphisms

Using a zero inflated linear model framework, we linked transcript-level variation in expression with single nucleotide alterations across the genome for the MOLM-13 dataset. Briefly, using genomic-coordinate intersection and the GenomicRanges R package, we first matched the genomic coordinates of SNVs with transcripts. Regarding the transcript coordinates, we used the Ensembl reported transcript start and transcript end to define the gene-body of a transcript and the 5000 bps upstream of the Ensembl reported transcription start site (TSS) to define potential cis-regulatory regions affecting that transcript. We created a matrix of expression and genotype locus (SNV) across all cells after determining the relevant SNV-Transcript pairings. Finally, utilizing this matrix, we fitted a zero-inflated linear model using the MAST R package with the following design: Transcript expression ∼ Genotype.

We used the GSEA-R tool in combination with the molecular signatures database (MSigDB) to examine enriched gene sets linked to differentially expressed genes across Molm-13 parental and resistant cells, as well as significant SNVs. In addition, using a default adjusted p-value of 0.10, we searched the Reactome Pathways database for meaningful pathways among these genes.

#### Significant variant testing

We created categorical variables for diploidy status and compared them using the chi-square test to identify differential SNVs between MOLM-13 P and R cells. Significant two-sided p-values were less than 0.05. Additionally, we used a multinomial logistic regression to determine if there were any variations in SNV prevalence between parental and resistant MOLM-13 types. We encoded the three states genotype (0/0, 0/1, 1/1) as the dependent variable and the MOLM-13 type (parental, resistant) as the independent variable for each SNP. A Wald Test was used to determine the model’s significance.

#### Sequencing Data Availability

DNA and RNA sequencing data is available for download in SRA (SRA ID to be added upon acceptance) for NA12878 and MOLM-13 cells. Sequencing data from the DCIS patient explored here is available through dbGAP (dbGAP ID to be added upon acceptance).

## ACKNOWLEDGEMENTS

We thank the patients involved in this study for their gracious gift of tissue samples that made the scientific insights in this manuscript possible.

## SUPPLEMENTARY DATA

**Supplementary Fig. 1.**
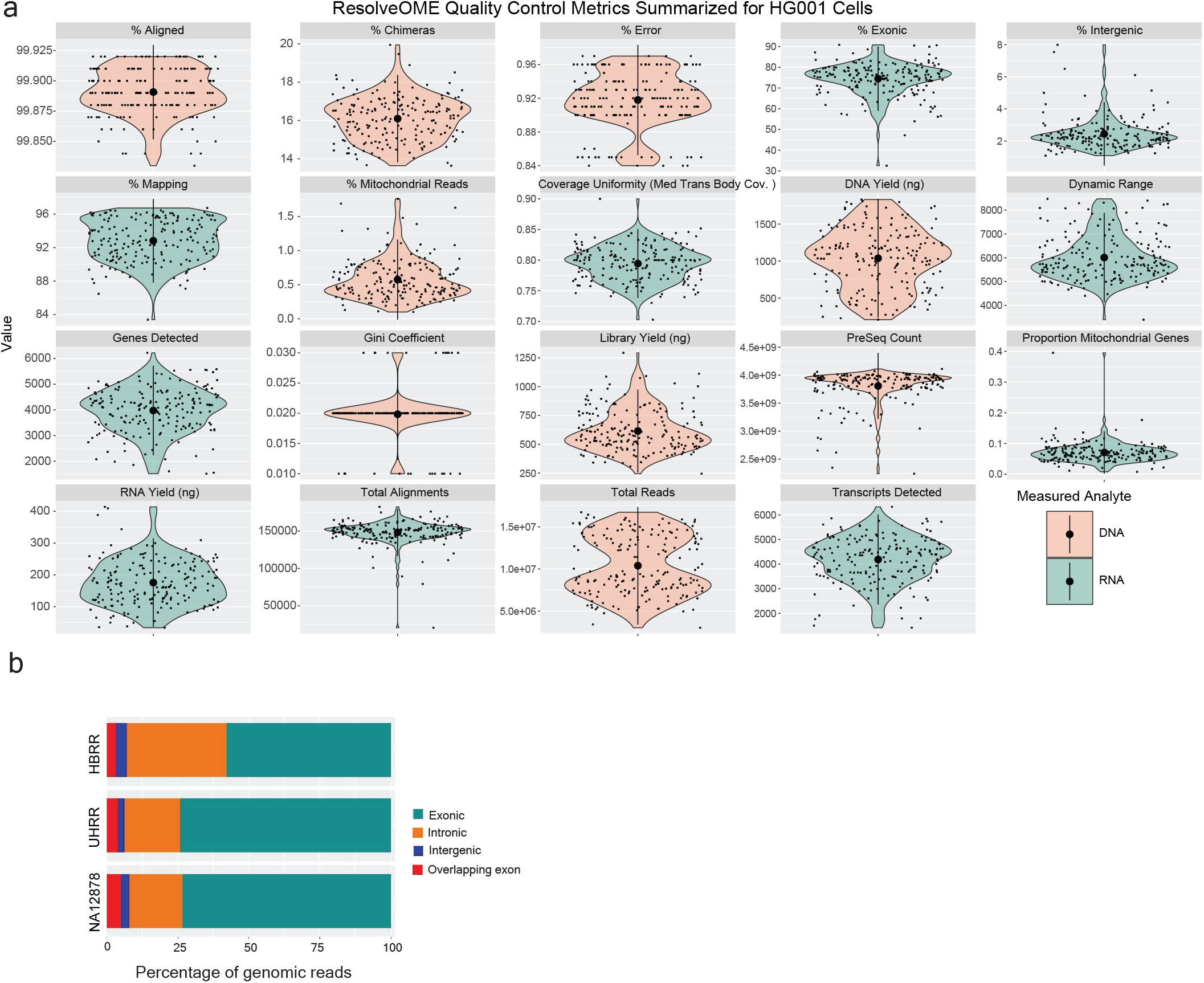
Performance metrics for genomic and transcriptomic arms of the workflow. **a,** Shown is mean +/- 2SD for the indicated metrics of single cell genomic (salmon) or transcriptomic (turquoise) ResolveOME libraries sequenced at low-depth. Single cells are denoted as points; shape of curve indicates overall distribution using the probability density function. The distribution of total reads for each cell is indicated for genomic libraries; transcriptomic libraries were down sampled to 100,000 total reads. **b**, Genomic feature mapping of single-cell NA12878 transcriptomes relative to Universal Human Reference RNA (UHRR) or Human Brain Reference RNA (HBRR) standards. n=165 (HBRR=28, UHRR=28, NA12878 = 109.

**Supplementary Fig. 2.**
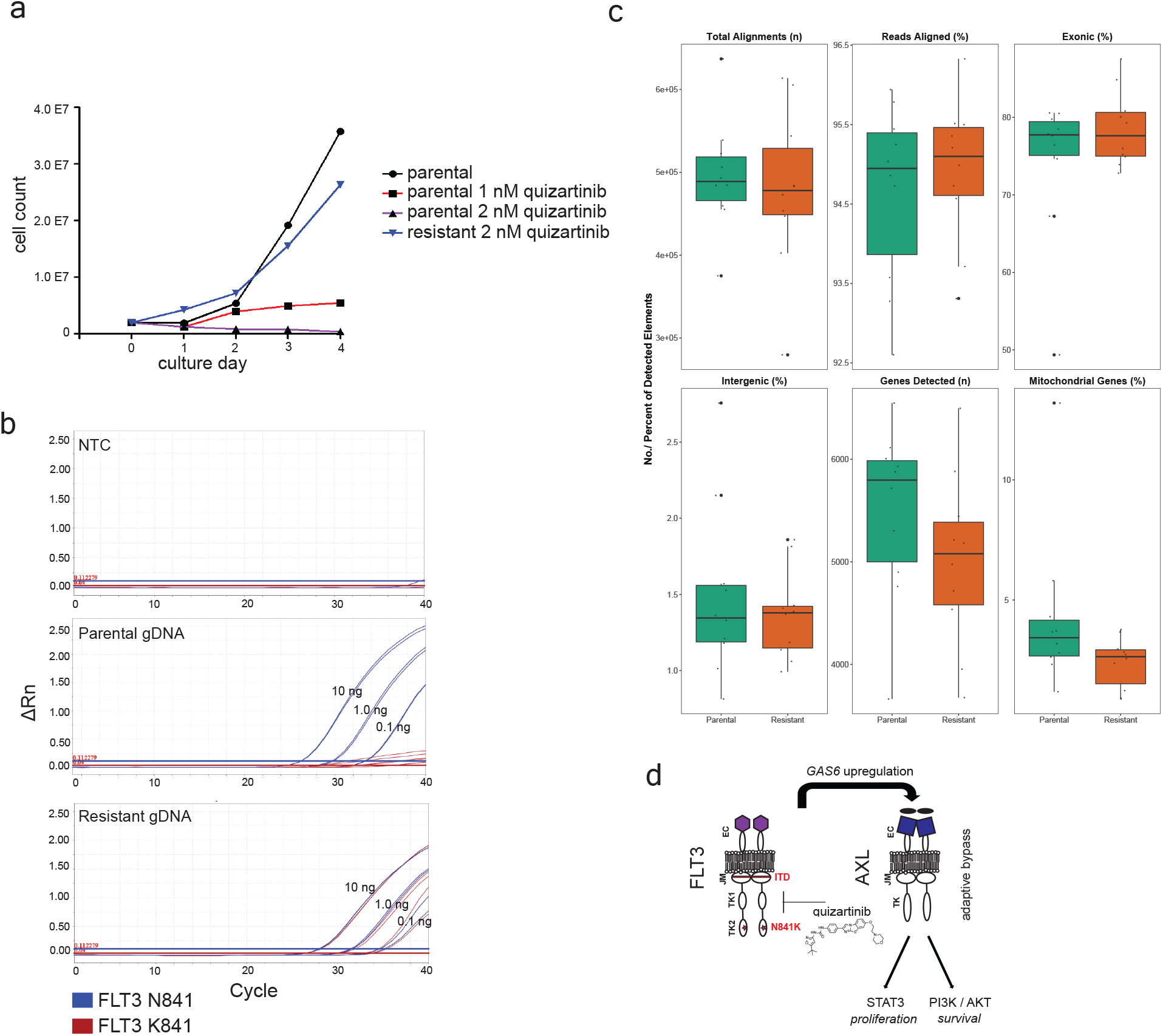
MOLM-13 AML quizartinib resistance model. **a**, Relative growth rates of parental and quizartinib-resistant MOLM-13 cells. Counts of cells over culture days after introduction of varying concentrations of quizartinib. **b**, qRT-PCR detection of mutant *FLT3* K841 in treatment-naïve parental cells. Cycling traces of *FLT3* N841 (blue) and K841 (red) in MOLM-13 parental and quizartinib-resistant cells. **c**, RNA performance metrics of parental and quizartinib-resistant MOLM-13 cells. Plots indicate number or percentage of each performance feature observed for a total of 20 single cells (Resistant = 10, Parental =10). **d**, Model of transcriptional bypass signaling through AXL upon FLT3 inhibition. Schematic illustrating that upon FLT3 inhibition by quizartinib, *GAS6*, a ligand for the receptor tyrosine kinase AXL, is transcriptionally upregulated in resistant MOLM-13 cells to drive growth and survival through PI3 kinase and AKT signaling, respectively.

**Supplementary Fig. 3.**
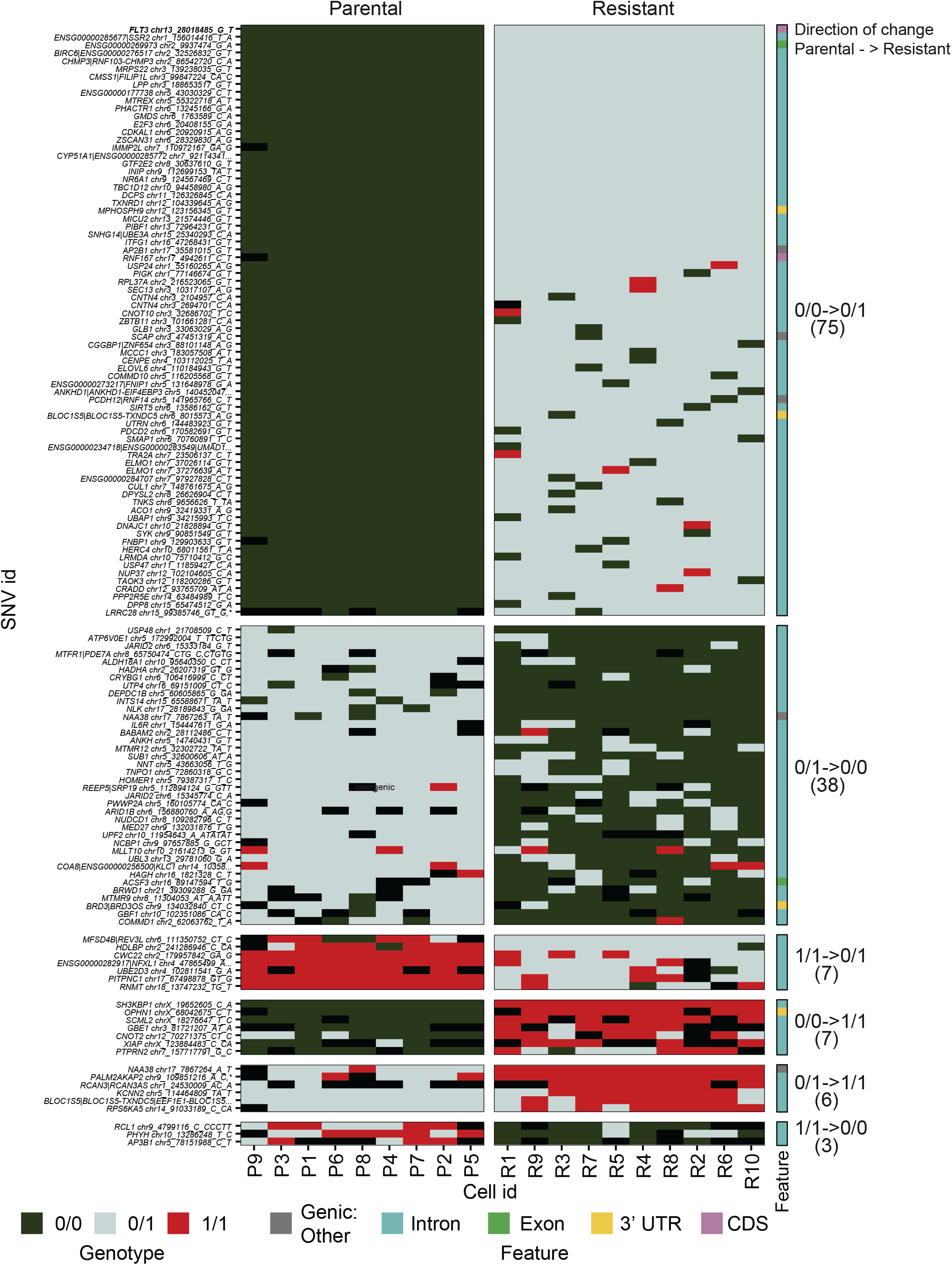
Single nucleotide variation between parental and quizartinib-resistant MOLM-13 cells. Selected significant single nucleotide variants (rows) with differential genotypic identity between parental and resistant cells (columns). *FLT3* N841K (chr13:28018485_G_T) is shown at the top of the heatmap. The number of cells residing in each category of Parental to Resistant direction of genotypic change is shown and the location of the SNV with regard to functional genomic element is shown (right bar, colors).

**Supplementary Table 1 | Variant annotation: MOLM-13 AML model of drug resistance.** Significant coding and non-coding variants in quizartinib-resistant vs. parental MOLM-13 cells. Positions that showed genotype association (Multinomial logistic-regression p-value) to parental vs. drug resistant are shown, but filtered for positions within or adjacent to coding sequence or for positions outside of coding space. Adjusted p-values are provided but were not filtered. File sorted for p-value and then by chromosome position. Cells were grouped according to genotype along with gene and transcript-level annotation is shown.

**Supplementary Table 2 | Differentially Expressed Transcripts between quizartinib-resistant and parental MOLM13-AML cells.** Differentially expressed transcripts/genes between parental and quizartinib resistant cells in the MOLM-13 model, estimated using a ZLM, fitted as a function of Cell Type (Parental/Resistant) and the Number of detected features (transcripts/genes) per cell. File sorted by adjusted p-values (ZLMpadj). P-values (ZLMpvalue) and fold change (ZLMFC) are also provided.

**Supplementary Table 3 | Differential Transcript Usage between quizartinib-resistant and parental MOLM13-AML cells.** Differential transcript usage analysis, performed between parental and resistant MOLM13 cells using a reconstructed matrix of transcript abundances based on scaled TPM. The transcript expression was modeled using the Dirichlet-multinomial distribution model. File includes the proportion of transcript A or B isoform in either resistant or parental cells, gene identity, and corresponding cluster shown in Figure 3a.

**Supplementary Table 4 | *CEBPA* upstream mutation coordinates.** We observed a genotypic bias in the frequency of *CEBPA* mutations between parental and resistant cells, characterized by specific locus and variant types.

**Supplementary Table 5| Ploidy:expression correlation in MOLM-13 AML model of drug resistance.** MOLM-13 genes with expression levels correlated with ploidy. A genome-ordered list of transcripts with shared expression magnitudes within known copy numbers from Figure 3b. Associated Gene IDs and Gene Symbols are provided.

**Supplementary Table 6| Regulatory variant discovery in MOLM-13 AML model of drug resistance: variant annotation and transcript expression values.** We identified single nucleotide variants (SNVs) associated with intracellular transcript expression in both resistant and parental cells. The results reveal candidate regulatory alleles and their corresponding transcripts that show differentiation between the groups depicted in Figure 3c. In addition to the transcript id, gene symbol, and genotype, the output includes raw z-score expression values and the proportion of each genotype within each group.

**Supplementary Table 7| Variant annotation: primary breast cancer biopsies.** Significant coding and non-coding variants in epithelial Normal vs Tumor cells. Positions that showed genotype association (Multinomial logistic-regression p-value) to Tumor vs. Normal are shown, but filtered for positions within or adjacent to coding sequence or for positions outside of coding space. Adjusted p-values are provided but were not filtered. File sorted for p-value and then by chromosome position.

**Supplementary Table 8| Gene expression count matrix for single cells from primary breast cancer biopsies.** The RNA abundance of ductal carcinoma in situ (DCIS) samples, quantified using the Salmon pseudo-alignment approach. The output provides expression summaries in the form of raw counts (countsFromAbundanceNo) and scaled counts (countsFromAbundancescaledTPM) represented as transcripts per million (TPM). The transcript-level abundance estimates generated by Salmon were mapped to corresponding genes using the R package tximport and aggregated to the gene-level by summing the transcript abundances for each gene.

**Supplementary Table 9| External datasets employed.** We pulled previously generated 10x data on NA12878 cell lines from a project in SRA (https://www.ncbi.nlm.nih.gov/bioproject/PRJNA521545). Using biosample SAMN10904684 (GEO: GSM3596321) we did UMI deconvolution to generate gene counts from ∼8000 cells. Additional SRA datasets employed in the manuscript are provided.

